# TCRMatch: Predicting T-cell receptor specificity based on sequence similarity to previously characterized receptors

**DOI:** 10.1101/2020.12.11.418426

**Authors:** William D. Chronister, Austin Crinklaw, Swapnil Mahajan, Randi Vita, Zeynep Kosaloglu-Yalcin, Zhen Yan, Jason A. Greenbaum, Leon E. Jessen, Morten Nielsen, Scott Christley, Lindsay G. Cowell, Alessandro Sette, Bjoern Peters

## Abstract

The adaptive immune system in vertebrates has evolved to recognize non-self-antigens, such as proteins expressed by infectious agents and mutated cancer cells. T cells play an important role in antigen recognition by expressing a diverse repertoire of antigen-specific receptors, which bind epitopes to mount targeted immune responses. Recent advances in high-throughput sequencing have enabled the routine generation of T-cell receptor (TCR) repertoire data. Identifying the specific epitopes targeted by different TCRs in these data would be valuable. To accomplish that, we took advantage of the ever-increasing number of TCRs with known epitope specificity curated in the Immune Epitope Database (IEDB) since 2004. We compared six metrics of sequence similarity to determine their power to predict if two TCRs have the same epitope specificity. We found that a comprehensive *k*-mer matching approach produced the best results, which we have implemented into TCRMatch, an openly accessible tool (http://tools.iedb.org/tcrmatch/) that takes TCR β-chain CDR3 sequences as an input, identifies TCRs with a match in the IEDB, and reports the specificity of each match. We anticipate that this tool will provide new insights into T cell responses captured in receptor repertoire and single cell sequencing experiments and will facilitate the development of new strategies for monitoring and treatment of infectious, allergic, and autoimmune diseases, as well as cancer.

## INTRODUCTION

T cells are lymphocytes that play a critical role in the function of the adaptive immune system [1]. Each T cell expresses a characteristic T-cell receptor (TCR) typically consisting of an α and β chain, which are formed during T cell maturation as a result of stochastic V(D)J gene recombination [2]. Different TCRs are capable of recognizing different epitopes presented by major histocompatibility (MHC) class I or class II proteins on the cell surface [3]. The specificity of a given TCR is dependent upon the amino acid sequence of each chain, particularly the three highly polymorphic complementarity-determining regions (CDR1, CDR2, CDR3). Broadly speaking, CDR3 directly interacts with the presented peptide, while CDR1 and CDR2 primarily interact with the MHC molecule [4]. The varying antigen specificity of different TCRs allows T cells to initiate immune responses against a broad and ever-changing range of non-self entities, including infectious agents and mutated cancer cells.

TCR repertoire sequencing has emerged as an accessible and efficient approach to capture the diversity of TCRs in blood or tissue samples of an individual [5]. Using different next-generation sequencing and bioinformatics approaches, TCRs can be sequenced to different levels of resolution, ranging from paired sequencing of full α and β chains to partial sequencing of the TCR β chain by itself. Single cell sequencing with targeted TCR identification, as provided by, for example, the 10x Genomics technology platform [6], identifies the gene expression state of individual T cells, along with the receptor sequences that – in principle – indicate the specificity of these T cells. All of the main TCR repertoire sequencing approaches in use today generate information about the CDR3 region in the TCR β chain, as that part of the TCR is thought to convey the most information about TCR specificity, and can serve as a “barcode” to track T cells with different specificities.

While TCR repertoire sequencing can track perturbations in the composition of antigen-specific T cell populations, it does not yield information on which TCRs are recognizing which epitopes. Such determination typically requires additional targeted experiments that can be challenging to perform. The desire to determine the specificity of T cells based on their receptor sequence has led to the development of different approaches. GLIPH and GLIPH2 [7, 8] were developed to cluster large sets of TCR sequences into groups with shared specificity, but these tools do not themselves predict likely recognized epitopes. Machine learning-based models designed to predict specificity of a receptor “*ab initio*” have also been developed [9, 10, 11]. These methods are computationally expensive and require large numbers of known TCRs and their epitopes for training, validation, and testing in order to avoid overfitting and learn models that generalize well. Thus, there is a need to develop new approaches to identify the specificity of TCRs in repertoire sequencing data that do not require additional experimentation, can recover the specificity for many different epitopes, and that are efficient enough to process current repertoire dataset sizes in reasonable times.

Here, we set out to address this challenge by taking advantage of the ever-growing dataset of epitopes in the Immune Epitope Database (IEDB) that have been experimentally determined to be recognized by T cells, and for which information on the specific TCR recognizing the epitope is also available [12, 13]. We tested several approaches to evaluate the sequence similarity between TCRs and examined how well they distinguished if the TCRs recognized the same epitopes or not. Using an initial test set of 24,678 TCR CDR3β sequences from the IEDB, we found that a comprehensive *k*-mer matching algorithm adopted from work by Shen *et al*. [14] performed best. This algorithm, which we called TCRMatch, also performed well on an independent dataset and has now been implemented as a web server (http://tools.iedb.org/tcrmatch/); it is also freely available for download as a standalone command-line tool.

## METHODS

### Compilation of TCR dataset

A dataset of CDR3β sequences and corresponding epitopes was compiled by querying the IEDB on May 30, 2020 for all curated TCR entries. Starting from the IEDB homepage (iedb.org), the following filters were applied: For Epitope, “Any Epitopes”; for Assay, “Positive Assays Only” and “T Cell Assays”; for Antigen, no filters for Organism or Antigen Name; for MHC Restriction, “Any MHC Restriction”; for Host, “Any Host”; and for Disease, “Any Disease”. Any records that lacked either a CDR3β sequence or at least one peptidic epitope were filtered out. CDR3β sequences were trimmed of excess flanking residues, where necessary, using a custom pHMM-based trimming tool (manuscript in preparation). This tool was based on the ImMunoGeneTics (IMGT) [15] notation of the CDR3β region, which excludes the constant N-terminal cysteine (C) residue and C-terminal phenylalanine (F) or tryptophan (W) residue. Following trimming, the dataset consisted of 24,973 receptor groups, each defined by a unique CDR3β sequence, and 993 unique peptidic epitopes. Of these 993 epitopes, 495 were recognized by only one receptor, which were excluded from the benchmarking analysis to ensure that all TCRs in the dataset had at least one other TCR recognizing the same epitope that could be identified. The final IEDB dataset consisted of 24,678 CDR3β sequences and 498 epitopes. Of the 24,678 receptor groups, 21,851 (88.5%) recognized one epitope, while 2,827 (11.5%) recognized multiple epitopes. As a control, a shuffled dataset was established that used the same CDR3β:epitope pairs from the final dataset, but shuffled the pairings between receptor groups and epitopes.

### 10x dataset

A dataset containing 15,769 CDR3β sequences and their epitope specificities was downloaded from a published application note by 10x Genomics [16]. Like the IEDB dataset, all 10x sequences were run through a custom trimming tool (manuscript in preparation) to ensure adherence to IMGT guidelines. Any CDR3β that did not recognize an epitope found in the IEDB was excluded in order to ensure successful predictions were possible for all sequences. Following this filtering step, the final dataset consisted of 3,218 CDR3β sequences recognizing one or more of 18 unique epitopes.

### Similarity metrics

Six scoring metrics were tested to measure similarity between all possible pairs of CDR3β sequences in the IEDB dataset (Table 1). We used Parasail [17] to carry out Needleman-Wunsch pairwise alignments using the BLOSUM62 substitution matrix. An open-gap penalty of −7 and extend-gap penalty of −1 were used in tandem with default scoring settings. From these alignments, we generated the metrics Identity Alignment, Identity Long, Identity Short, and Alignment Score. “Identity” metrics were calculated by dividing the number of exact matches by either the length of the overlap between the aligned sequences (Identity Alignment), the length of the longer sequence, where applicable (Identity Long), or the length of the shorter sequence, where applicable (Identity Short). Alignment Score was calculated by dividing the score of the optimal alignment by the length of the alignment overlap. We also increased the Parasail gap penalties to −50 and −20 (open- and extend-gap, respectively) to test whether precision and recall improved when gaps were reduced for each of the four aforementioned metrics; however, we did not find any significant change in performance. Levenshtein distance, also known as edit distance, was measured between sequences using an open source Python implementation of the algorithm (https://pypi.org/project/python-Levenshtein). TCRMatch was implemented from work by Shen *et al*. [14].

**Table 1:** Description of the scoring metrics used to identify receptors with identical epitopes.

| Metric | Description |
| --- | --- |
| Alignment Score | Alignment score divided by length of alignment |
| Identity Alignment | Percent identity within length of alignment |
| Identity Long | Percent identity within length of longer sequence |
| Identity Short | Percent identity within length of shorter sequence |
| Levenshtein | Minimum number of edits (substitutions, insertions, and deletions) necessary to transform one sequence into another |
| TCRMatch | Comprehensive comparison of all possible $k$-mers using BLOSUM62 observed frequency matrix |

The TCRMatch algorithm relies on a modified version of the BLOSUM62 observed frequency matrix for amino acid substitutions [14]. The two sequences being tested for similarity, seq1 and seq2, are split into sets of *k*-mers, beginning with *k*=1 and incrementing by 1 up to the length of the sequences, or the shorter sequence if seq1 and seq2 differ in length. For each value of *k*, all possible combinations of *k*-mer pairs between the seq1 set and seq2 set are compared and assigned a similarity value derived from the values of the transformed BLOSUM62 matrix. For *k* = 1, values for each amino acid pair are looked up and summed. For *k* > 1, each amino acid of each seq1 *k*-mer is compared to the amino acid at the same position in the seq2 *k*-mer, and the matrix values are multiplied together prior to being summed across all possible *k*-mer pairs. The sums from each value of *k* are added together and divided by the normalization factor, 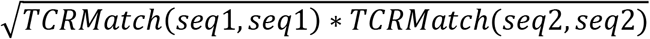, to yield a score between 0 and 1, where 1 signifies a perfect match (i.e., seq1 = seq2).

### Evaluation of prediction performance using precision and recall

To test a given similarity metric, it was applied to compare each of the 24,678 CDR3β sequences to all other CDR3β sequences in the same dataset. A range of similarity cutoffs were employed for each metric, and CDR3β sequences surpassing the similarity cutoff (“matches”) were considered true positives (TPs) or false positives (FPs) based on whether the matching CDRs recognized the same epitope or not. A TP was defined as an epitope recognized by the match sequence that was also recognized by the input CDR3β. Conversely, a false positive (FP) was defined as a match epitope that was not recognized by the input CDR3β. For a given similarity cutoff, precision was defined as TP/(TP+FP), while recall was defined as the fraction of epitopes recognized by input CDR3β sequences for which a matching CDR3β sequence with the same epitope was identified.

### Bootstrapping approach to compare metrics

To compare the performance of similarity metrics, we simulated 100 datasets by sampling (with replacement) 24,678 CDR3β sequences from the IEDB dataset. Each sampled sequence was scored for similarity to the remaining 24,677 sequences using the algorithms Identity Long, TCRMatch, and Alignment Score. Precision and recall were calculated at each score threshold for each method. For each precision-recall curve, we calculated the area-under-the-curve (AUC) for the recall range of 0 to 0.5, yielding a distribution of 100 AUCs for each method. The AUCs of each method were compared in pairwise fashion, and the fraction of the 100 comparisons in which the AUC of one method exceeded the AUC of the other (i.e., the p-value) was determined.

## RESULTS

### Assembly of TCR dataset with known epitopes for cross-validation from the IEDB

We used the IEDB [12] to retrieve T cell receptors and their known epitope specificity to serve as our benchmark dataset (see Methods). Receptors were assembled into groups defined by a unique CDR3β sequence found in the IEDB that had one or more corresponding epitopes. Our goal was to evaluate algorithms that take a query CDR3β as an input, identify similar CDR3β sequences in the database, and determine how well the similarity in CDR3B predicts that different TCRs recognize the same epitope. Given that goal, we excluded epitopes for which only one CDR3β existed. All told, our IEDB dataset contained 24,678 receptor groups, each with specificity for one or more known epitopes.

### Selection of similarity metrics

We set out to identify established sequence similarity metrics that would be most applicable for comparing TCR sequences (Table 1). We identified four metrics based on pairwise alignment of CDR3β sequences, one on edit distance, and one on *k*-mer composition. Alignment-based methods are an established approach to measuring similarity between amino acid sequences, including TCRs [19]. We carried out Needleman-Wunsch pairwise alignments for all sequence pairs and computed four similarity scores: Alignment Score, Identity Alignment, Identity Long, and Identity Short. Similarly, Levenshtein distance, which has been used previously for similarity-based clustering of TCR sequences [20], was calculated for each pair of TCR sequences. Finally, the TCRMatch algorithm was implemented based on the work of Shen *et al*. [14] and used to calculate similarity scores for all IEDB pairs. This algorithm has been shown to be effective in identifying MAIT cell TCRs from unknown sequences [18].

### Establishment of performance evaluation metrics

Each of the TCR sequence similarity metrics was tested for its ability to take input TCR sequences with known epitopes and match them with similar TCRs that recognize the same epitope. To measure performance, we calculated precision and recall on the epitopes recovered from such matches in cross-validation for a series of similarity cutoffs (Fig. 1).

**Figure 1:**
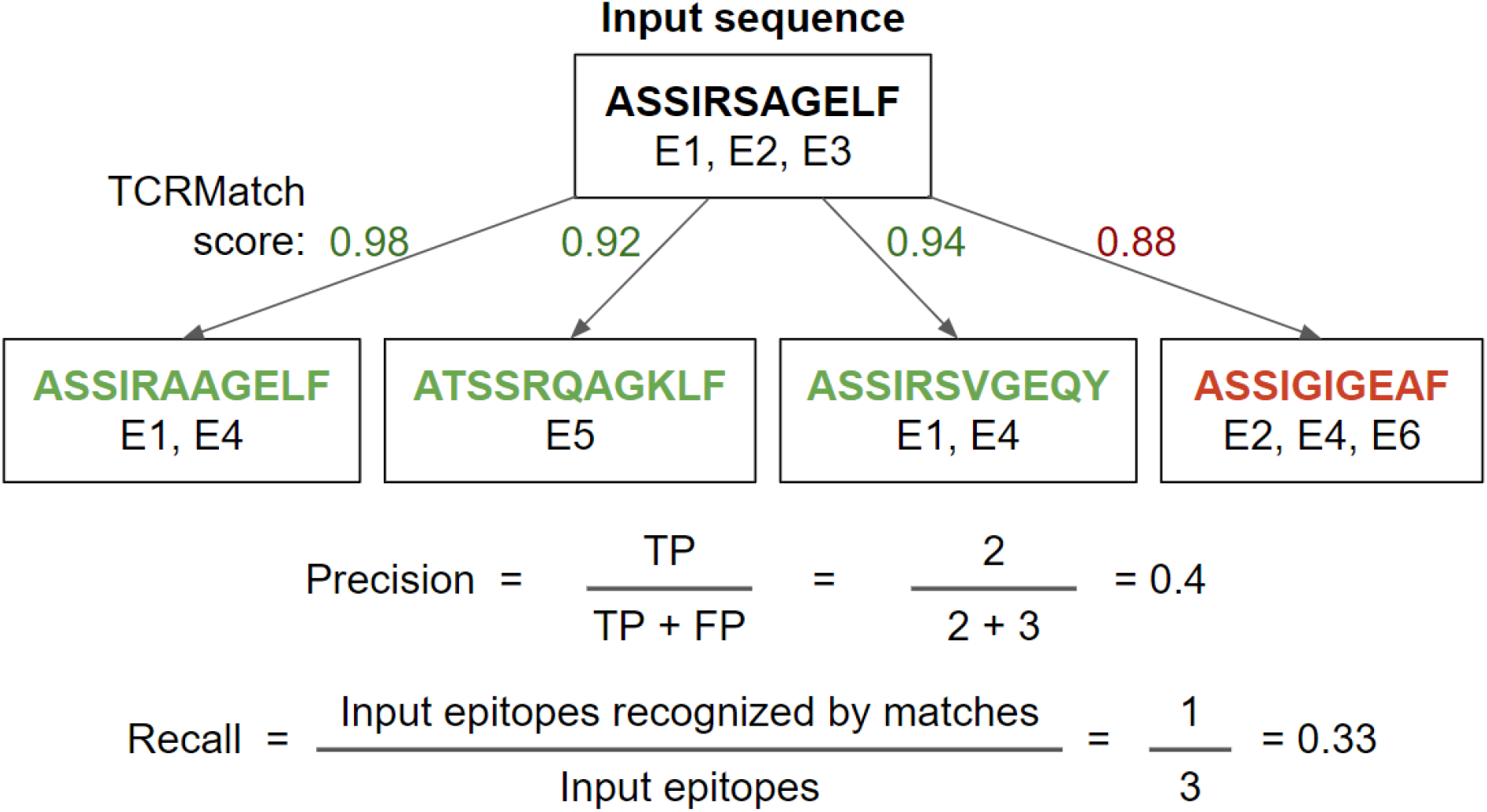
Example precision and recall calculation. Using a TCRMatch score cutoff of 0.9, the input sequence matches 3 out of 4 tested sequences. Of the 5 epitopes recognized by the 3 matches, 2 epitopes (E1, E1) are shared with A (true positives, TP) and 3 epitopes (E4, E5, E4) are not shared with A (false positives, FP), resulting in a precision of 2/5. Of the input sequence’s three epitopes (E1, E2, E3), E1 is also recognized by the first and third match, while E2 and E3 are not recognized by any of the matches; therefore, recall = 1/3.

### Comparison of algorithms on IEDB dataset and randomized dataset

We tested six similarity metrics on IEDB data and evaluated the relative success of each method using precision and recall (Fig. 2). TCRMatch was the strongest performer for recall values between 0.05 and 0.40. For example, at a similarity cutoff of 0.94, TCRMatch showed a recall of 0.213 and a precision of 0.532; in other words, for 21.3% of epitopes recognized by input TCRs, TCRMatch correctly identified a matching TCR recognizing the same epitope (recall), and 53.2% of the total match calls made at that threshold were correct (positive predictive value). At similar recall levels, ranging from 0.196 to 0.230, Levenshtein distance, Alignment Score, Identity Long, and Identity Alignment yielded maximum precision rates ranging between 0.441 and 0.478. The poorest performing metric was Identity Short, which at a comparable recall of 0.255 yielded a precision of 0.113; moreover, across all thresholds tested, Identity Short never exceeded 0.124 in precision. These subpar results were largely driven by perfect or near-perfect alignments of short CDR3β sequences to longer CDR3β sequences, which proved to be only weakly associated with finding a matching epitope. Starting around a recall of 0.5 and precision of 0.25, all methods except for Identity Short began to show similar precision as recall increased towards 1.

**Figure 2:**
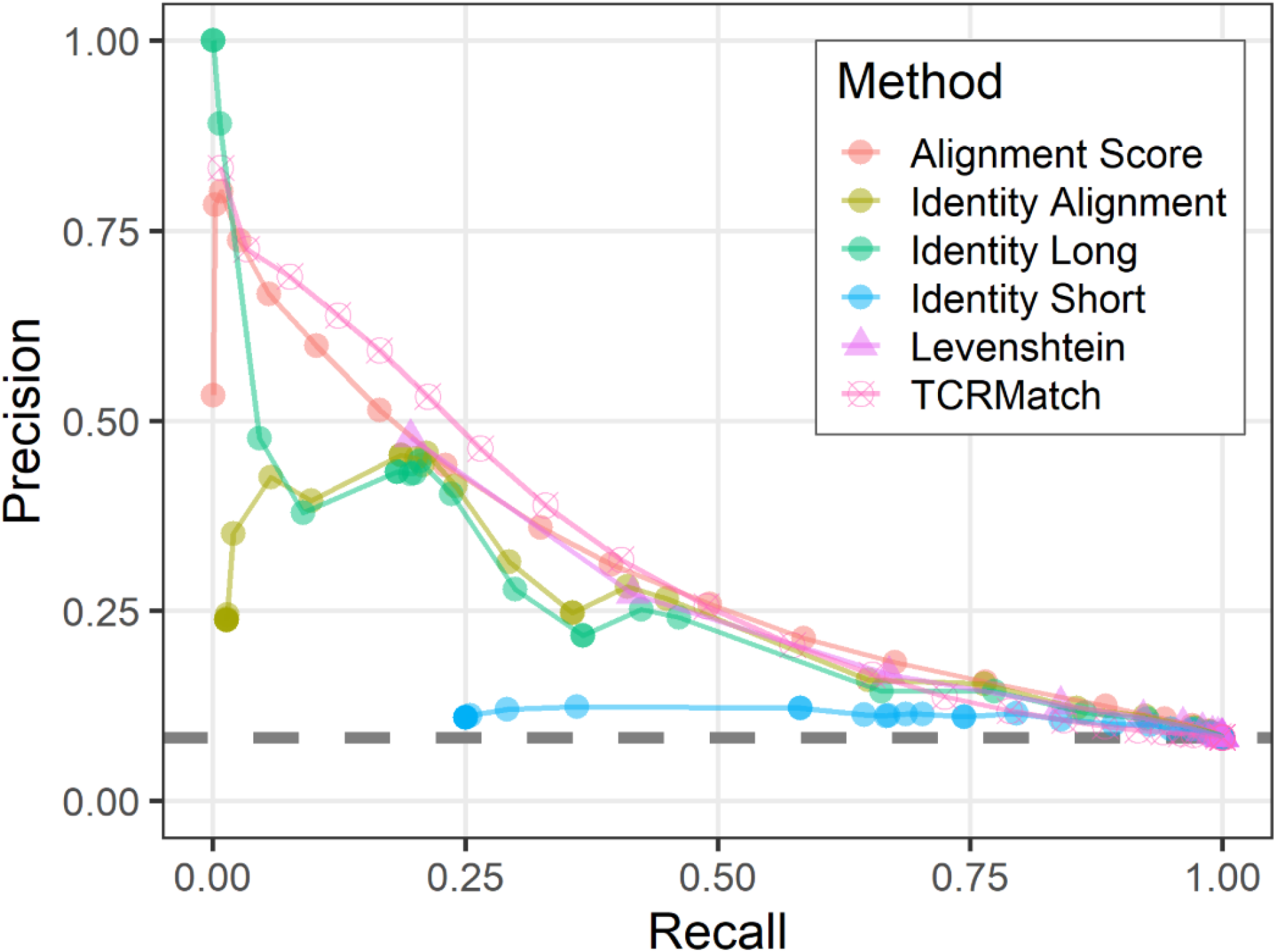
Precision-recall plots comparing different sequence similarity metrics. Each similarity metric was evaluated at different thresholds for its ability to recall TCR sequences in the IEDB that recognized the same epitope (x-axis) and compared that to the precision at the same threshold (y-axis), which specifies the percentage of match epitopes that were also recognized by the input sequences. The gray dashed line indicates average performance on the randomized IEDB dataset, wherein CDR3β-epitope pairs were shuffled.

Upon inspecting the precision-recall graph, it was apparent that there were three metrics that had a precision above all others at a given recall: Identity Long (for the recall range of 0 to 0.02), TCRMatch (0.02 to 0.47), and Alignment Score (0.47 to 0.75). To compare these three metrics statistically, we performed a bootstrapping analysis (see Methods) to calculate the area-under-the-curve (AUC) for the part of the graph where precision values were at least ~3 times higher than the baseline observed in analyzing a randomized dataset, (baseline precision = 0.084, dashed line in Fig 2). This corresponded to recall values between 0.0 and 0.5. This bootstrapped AUC analysis showed that the TCRMatch AUC of 0.255 was significantly higher than the AUC of Identity Long, 0.203 (p < 0.01), and Alignment Score, 0.245 (p < 0.01).

Analysis of the randomized dataset showed precision consistently fluctuating around 0.084 regardless of similarity metric used, a value consistent with the precision found when analyzing the non-randomized dataset without any similarity thresholds imposed. This result demonstrates that all of the tested similarity metrics are better than a randomized control at identifying receptors with the same epitope specificity, and that the best performing methods far exceed the random performance, especially at more stringent thresholds.

### Performance evaluation on independent test dataset

Following initial tests based on the IEDB dataset, which showed TCRMatch to be the highest performing metric, we assessed its performance on an independent, publicly available dataset from 10x Genomics [16]. This dataset was generated using TCR repertoire sequencing of T cells found to bind to pMHC multimers presenting peptides from various viral and cancer proteins. Initially, this dataset contained 15,769 CDR3β sequences recognizing a total of 39 epitopes. The CDR3β sequences were trimmed of any flanking residues in accordance with IMGT notation [15] using a custom pHMM-based tool (manuscript in preparation) prior to analysis. To utilize this dataset for performance evaluation of a classifier trained on IEDB data, we used the subset of the 10x TCRs that recognized one of the 18 epitopes also recognized in the IEDB, resulting in a total dataset of 3,218 CDR3β sequences.

We used the 10x dataset as input and asked if TCRs from this dataset could have been assigned a putative epitope by searching for similar TCRs in the IEDB dataset. For this analysis, we used the similarity metrics TCRMatch, the top performer from the IEDB dataset according to the bootstrapped AUC analysis, and Levenshtein distance, another strong performer that was not part of the AUC analysis (Fig. 2). We observed similar results to those of the original dataset; TCRMatch outperforms Levenshtein in precision when recall is below 0.50, but the methods begin to converge as recall exceeds 0.50 (Fig. 3). The precision and recall of both metrics benefited from 232 10x sequences that had exact matches in the IEDB; of these matches, 171 (73.7%) recognized the same epitope. Overall, precision was lower than when the IEDB dataset was tested against itself (Fig. 2), as expected.

**Figure 3:**
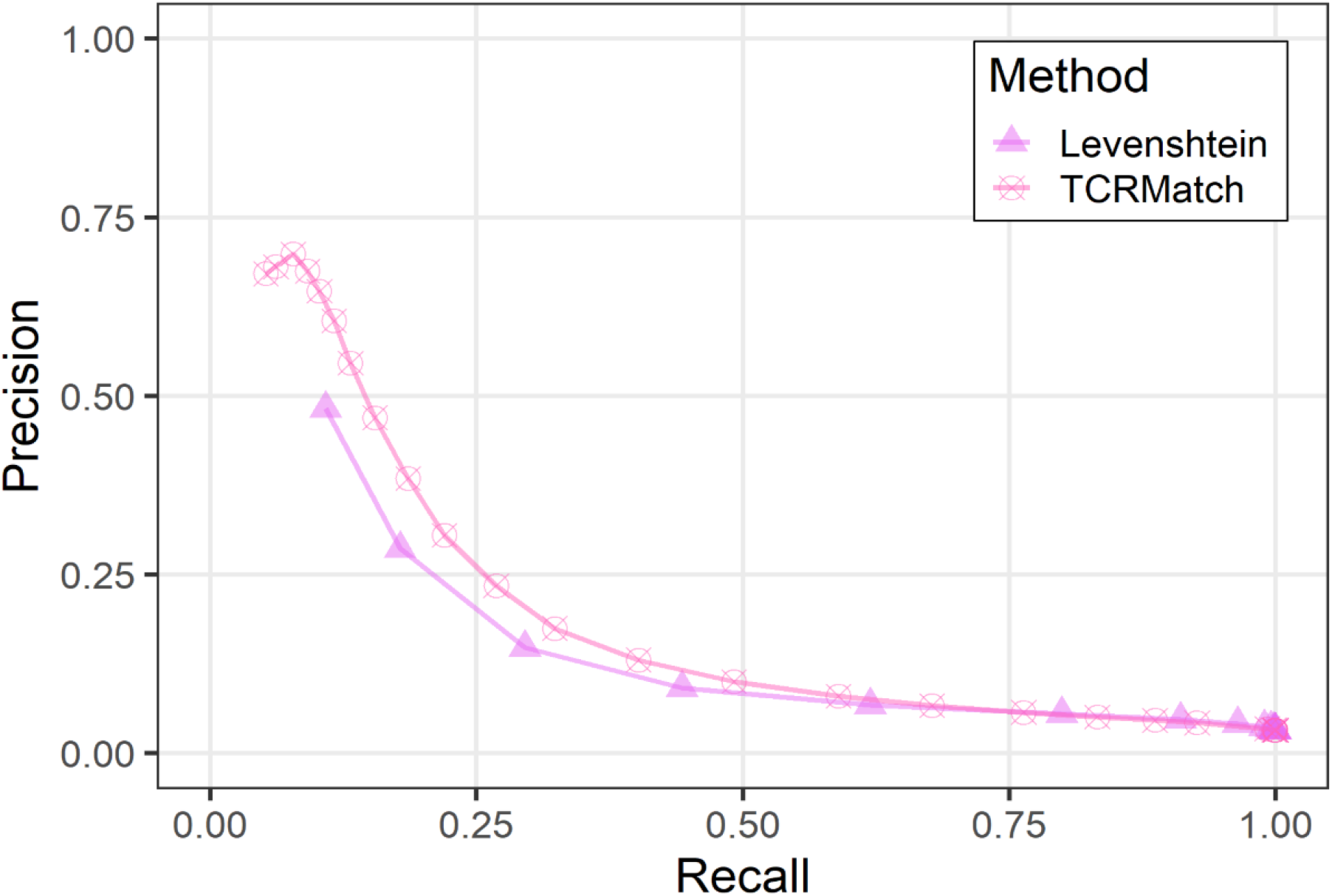
TCRMatch outperforms Levenshtein distance in 10x dataset. Comparison of TCRMatch and Levenshtein precision and recall metrics across similarity thresholds in analysis of 10x dataset against IEDB.

### Implementation of a Web Server

The motivating use case for our work has been to enable users to determine what the likely epitopes recognized are for their TCR(s) of interest. We have implemented TCRMatch as a web-based tool hosted on the IEDB (Fig. 4). A user can upload a set of up to 500 CDR3β sequences to query against the IEDB. Stringency can be specified by the user through the score threshold parameter. The TCRMatch algorithm is then run on the back end and results exceeding the user’s score cutoff are returned in tabular format, which can be downloaded as a CSV file. For datasets exceeding 500 sequences, TCRMatch analysis is also available through VDJServer (vdjserver.org) after creating a free VDJServer account [21]. Additionally, users have the option to download and run a standalone version of TCRMatch (github.com/IEDB/TCRMatch) to analyze large datasets.

**Figure 4:**
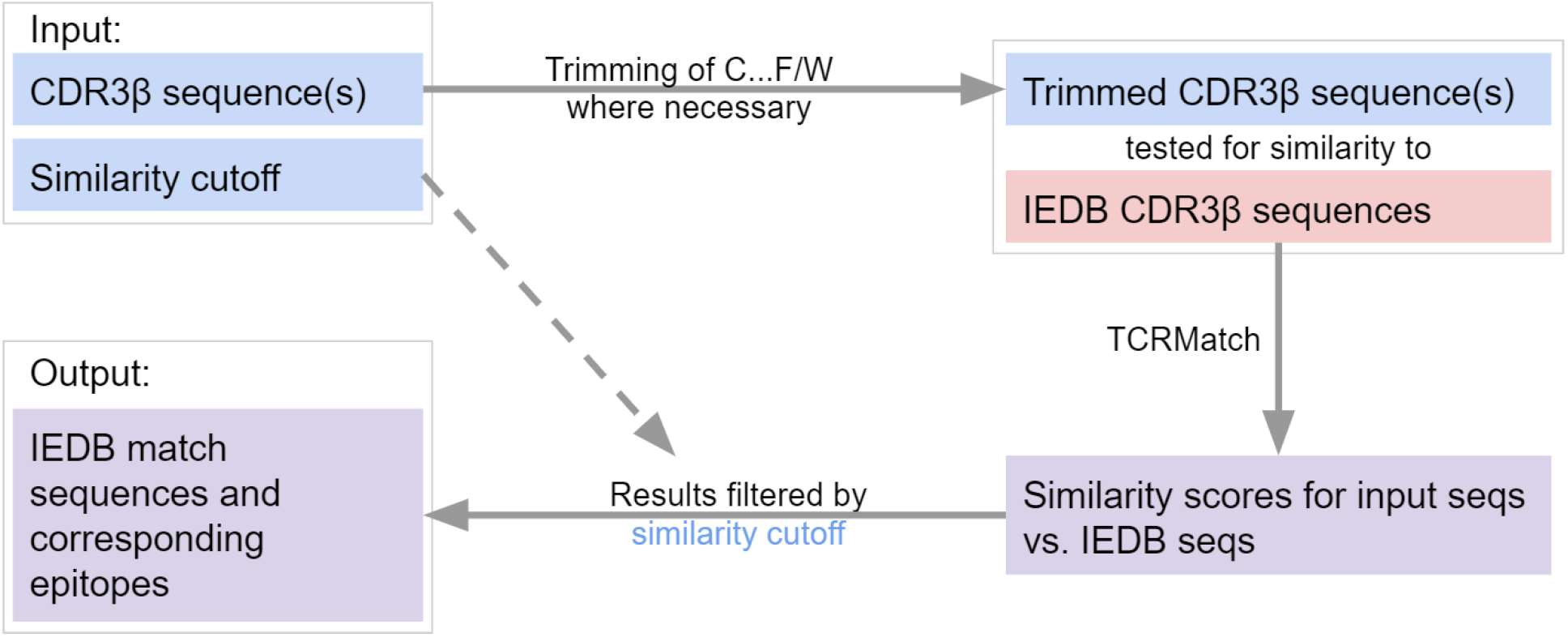
Flowchart of TCRMatch. The user provides one or more CDR3β sequences and selects a similarity cutoff. If N-terminal cysteine (C) and C-terminal phenylalanine (F) or tryptophan (W) are present, these residues are removed prior to the similarity search against the IEDB CDR3β sequences. The chosen similarity cutoff is used to filter TCRMatch’s final results, which consist of matching sequences and corresponding epitopes from the IEDB.

## DISCUSSION

T cells recognize epitopes presented to them by MHC molecules. Methods to identify epitopes recognized by T cells have primarily focused on the prediction of epitope binding to MHC molecules, which has been a cornerstone of immuno-informatics [22, 23]. More recently, methods analyzing T-cell repertoire sequencing data have been derived, to determine the specificity of a T cell based on its TCR sequence [9, 10, 11]. Such methods are often computationally expensive and offer limited predictive power due to the data on which they were trained. While we anticipate a prominent role for such approaches when the critical mass of data becomes available, until then, approaches that can efficiently characterize the hundreds of thousands of sequences generated by TCR repertoire sequencing without requiring a trained model represent a straightforward solution to an unmet need. To meet this need, we introduce TCRMatch, a method that balances efficiency and predictive power to find relevant TCRs through sequence similarity. Our method utilizes the power of *k*-mer-based methods for computing sequence similarity with a low computational footprint. This approach allows for the processing of full repertoire sequencing datasets in an efficient and accurate manner.

The method was tested on the large amount of TCR sequence data available through the IEDB, as well as on an independent test repertoire sequencing dataset published by 10x Genomics [16]. These analyses supplied evidence that the *k*-mer-based approach of TCRMatch provides a significantly higher number of relevant results compared to alignment-based or edit distance-based methods, in addition to being computationally efficient.

As our tool is built around the CDR3β sequences stored in the IEDB, the ability of TCRMatch to find matches is limited by the number and diversity of curated sequences available in the database. The baseline precision of 0.084 for the randomized control dataset (Fig. 2) indicates that the TCR-epitope data published to date, and collected by the IEDB, is biased towards a subpopulation of well-characterized epitopes. If the IEDB dataset containing 498 unique epitopes had an equal number of TCRs for each epitope, the expected baseline precision would have been approximately 1/498 (0.002). Instead, the frequency of TCRs per epitope varied considerably, with the five most common epitopes accounting for 15,243 of 28,001 (54.4%) total epitopes recognized. Studies of less characterized or uncharacterized epitopes will be beneficial for expanding the capability of TCRMatch to find matches for input CDR3β sequences. Nonetheless, it is important to note that the imbalance of epitopes in curated TCR data has no effect on the accuracy of the tool in finding matching sequences. Furthermore, researchers interested in searching CDR3β sequences against custom databases can download and modify the TCRMatch code (github.com/IEDB/TCRMatch) to do so.

Currently, TCRMatch identifies matches based solely on the input CDR3β sequence, which is the most closely associated with TCR specificity. As more data becomes available, we anticipate that a more complex version of TCRMatch, integrating additional data such as CDR3α sequences, CDR1 and CDR2 sequences from both α and β chains, and MHC restriction data, will become available and improve the accuracy of the tool’s epitope predictions.

The TCRMatch tool we develop here is well suited to complement existing tools such as GLIPH2 [8], which identifies clusters of TCRs in experimental data that likely recognize the same epitope. For example, a researcher who carries out TCR repertoire sequencing while studying a particular immune response might use GLIPH2 to discover a few hundred receptor clusters; each of those clusters can then be queried for their putative epitopes using TCRMatch, which further can enable follow-up experiments studying the epitopes driving the immune response in question. As the IEDB continues to accumulate immunological data, the quality and quantity of results produced by TCRMatch will continue to improve.

## REFERENCES

1. Buchholz, V. R., Schumacher, T. N. M. & Busch, D. H. T Cell Fate at the Single-Cell Level. Annu. Rev. Immunol. 34, 65–92 (2016).

2. Bradley, P. & Thomas, P. G. Using T Cell Receptor Repertoires to Understand the Principles of Adaptive Immune Recognition. Annu. Rev. Immunol. 37, 547–570 (2019).

3. Rudolph, M. G., Stanfield, R. L. & Wilson, I. A. HOW TCRS BIND MHCS, PEPTIDES, AND CORECEPTORS. Annu. Rev. Immunol. 24, 419–466 (2006).

4. Rossjohn, J. et al. T Cell Antigen Receptor Recognition of Antigen-Presenting Molecules. Annu. Rev. Immunol. 33, 169–200 (2015).

5. Calis, J. J.A. & Rosenberg, B. R. Characterizing immune repertoires by high throughput sequencing: strategies and applications. Trends in Immunology 35, 581–590 (2014).

6. Zheng, G. X. Y. et al. Massively parallel digital transcriptional profiling of single cells. Nature Communications 8, 14049 (2017).

7. Glanville, J. et al. Identifying specificity groups in the T cell receptor repertoire. Nature 547, 94–98 (2017).

8. Huang, H., Wang, C., Rubelt, F., Scriba, T. J. & Davis, M. M. Analyzing the Mycobacterium tuberculosis immune response by T-cell receptor clustering with GLIPH2 and genome-wide antigen screening. Nature Biotechnology 1–9 (2020). doi:10.1038/s41587-020-0505-4

9. Gielis, S. et al. Detection of Enriched T Cell Epitope Specificity in Full T Cell Receptor Sequence Repertoires. Front. Immunol. 10, 2820 (2019).

10. Fischer, D. S., Wu, Y., Schubert, B. & Theis, F. J. Predicting antigen specificity of single T cells based on TCR CDR3 regions. Molecular Systems Biology 16, e9416 (2020).

11. Dash, P. et al. Quantifiable predictive features define epitope-specific T cell receptor repertoires. Nature 547, 89–93 (2017).

12. Vita, R. et al. The Immune Epitope Database (IEDB): 2018 update. Nucleic Acids Res. 47, D339–D343 (2019).

13. Mahajan, S. et al. Epitope Specific Antibodies and T Cell Receptors in the Immune Epitope Database. Front Immunol 9, 2688 (2018).

14. Shen, W.-J., Wong, H.-S., Xiao, Q.-W., Guo, X. & Smale, S. Towards a Mathematical Foundation of Immunology and Amino Acid Chains. arXiv:1205.6031 [cs, q-bio, stat] (2012).

15. Lefranc, M.-P. et al. IMGT®, the international ImMunoGeneTics information system® 25 years on. Nucleic Acids Res 43, D413–D422 (2015).

16. A New Way of Exploring Immunity—Linking Highly Multiplexed Antigen Recognition to Immune Repertoire and Phenotype. 1–13 (10x Genomics, 2020).

17. Daily, J. Parasail: SIMD C library for global, semi-global, and local pairwise sequence alignments. BMC Bioinformatics 17, 81 (2016).

18. Wong, E. B. et al. TRAV1-2+ CD8+ T-cells including oligoconal expansions of MAIT cells are enriched in the airways in human tuberculosis. Commun Biol 2, 203 (2019).

19. Meysman, P. et al. On the viability of unsupervised T-cell receptor sequence clustering for epitope preference. Bioinformatics 35, 1461–1468 (2019).

20. Friedensohn, S., Khan, T. A. & Reddy, S. T. Advanced Methodologies in High-Throughput Sequencing of Immune Repertoires. Trends in Biotechnology 35, 203–214 (2017).

21. Christley, S. et al. VDJServer: A Cloud-Based Analysis Portal and Data Commons for Immune Repertoire Sequences and Rearrangements. Front Immunol 9, 976 (2018).

22. Peters, B., Nielsen, M. & Sette, A. T Cell Epitope Predictions. Annu. Rev. Immunol. 38, 123–145 (2020).

23. Dhanda, S. K. et al. IEDB-AR: immune epitope database—analysis resource in 2019. Nucleic Acids Research 47, W502–W506 (2019)

